# Decreased IgG core fucosylation, a player in antibody-dependent cell-mediated cytotoxicity, is associated with autoimmune thyroid diseases

**DOI:** 10.1101/362004

**Authors:** Tiphaine C. Martin, Mirna Šimurina, Marta Ząbczyńska, Marina Martinić Kavur, Magdalena Rydlewska, Marija Pezer, Kamila Kozłowska, Andrea Burri, Marija Vilaj, Renata Turek-Jabrocka, Milena Krnjajić-Tadijanović, Małgorzata Trofimiuk-Müldner, Anna Lityńska, Alicja Hubalewska-Dydejczyk, Irena Trbojević-Akmačić, Ee Mun Lim, John P. Walsh, Ewa Pochec, Tim D. Spector, Scott G. Wilson, Gordan Lauc

## Abstract

Autoimmune thyroid diseases (AITD) are the most common group of autoimmune diseases, associated with lymphocyte infiltration and the production of thyroid autoantibodies, like thyroid peroxidase antibodies (TPOAb), in the thyroid gland. Immunoglobulins (Igs) and cell-surface receptors are glycoproteins with distinctive glycosylation patterns that play a structural role in maintaining and modulating their functions. We investigated associations of total circulating IgG and peripheral blood mononuclear cells (PBMCs) glycosylation with AITD and the influence of genetic background. The study revealed an inverse association of IgG core fucosylation with TPOAb and PBMCs antennary α1,2 fucosylation with AITD, but no shared genetic variance between AITD and glycosylation. These data suggest that the decreased level of IgG core fucosylation is a risk factor for AITD that promotes antibody-dependent cell-mediated cytotoxicity (ADCC) associated with TPOAb levels.

## Introduction

Autoimmune thyroid diseases (AITD) are a class of chronic, organ-specific autoimmune disorders that disturb the function of the thyroid gland. They affect close to 5% of the European population (with a gender disparity) and so, represent the most common group of autoimmune diseases^1^. AITD encompass a spectrum of conditions including Hashimoto’s thyroiditis (HT) and Graves’ disease (GD). One of the features of AITD is the production of autoantibodies against components of thyroid cells that are also detected in the bloodstream.

The autoantibodies are produced against the three core thyroid proteins: thyroid peroxidase (TPO), thyroglobulin (Tg), and the thyroid-stimulating hormone receptor (TSH-R). Except for antibodies against TSH receptors (TSAb), which are known to stimulate the production of thyroid hormones by binding TSH receptors in GD^2^, little is known about the role of two other thyroid autoantibodies, thyroid peroxidase antibodies (TPOAb) and thyroglobulin antibodies (TgAb). Circulating TPOAb is the most common and diagnostically useful marker of AITD, detectable in the serum of most HT (95%) and GD (85%) patients^3^. In recent years, using TPOAb as a marker has been challenged since it appears in approximately 10% of apparently healthy individuals^4^. Even though autoantibodies are often a hallmark of autoimmune disorders, they can appear years before the first symptoms^5^, which poses the question about their causative role. Some evidence exists that autoantibodies can trigger autoimmunity, and IgG isotype seems to be connected with the development of autoimmune diseases potentially through regulation of IgG effector functions by alternative glycosylation^5^. However, it is yet to be determined if anti-thyroid antibodies cause AITD, or whether additional control mechanisms, such as post-translational modifications, are required to trigger the disease onset.

The most abundant and diverse form of post-translational modification is glycosylation, the attachment of sugar moieties to proteins, and various glycans are involved in virtually all physiological processes^6^. Glycans attached to Immunoglobulin G (IgG) are indispensable for its effector function and control of inflammation^7-10^. There are two glycosylation sites within the fragment crystallizable (Fc) of IgG that affect the molecule’s 3D-conformation and affinity for binding to Fcγ-receptors (FcγRs) on a range of immune cells^6,11,12^. Additional N-glycosylation sites are present in approximately 20% of IgG fragment antigen binding (Fab) and play a role in immunity, such as the affinity of epitope-binding site^13-16^. Previous analysis of IgG glycosylation with other autoimmune diseases showed a reduction of IgG galactosylation and sialylation, which trigger inflammatory response^17-20^. In relation to AITD, two small studies (62 and 146 patients respectively) looked at the TgAb glycosylation and reported differences between AITD, papillary thyroid cancers (PTC), and controls^21,22^. First, TgAb IgG from HT patients showed higher levels of core fucose than the control group, as well as of terminal sialic acid and mannose^21^. On the other hand, it was observed that among HT, GD, and PTC groups, HT patients had significantly lower core fucose content on TgAb than the other two groups^22^. Furthermore, since recent genome-wide association studies (GWAS) identified novel loci associated with IgG glycosylation, which were known to be strongly associated with autoimmune conditions^23,24^ and the heritability of AITD is estimated to 55-75%^25-27^, the next logical step was to examine if there was any common genetic background between those features. No data is currently available on common genetic variants associated with IgG glycosylation traits and AITD, and no large study on associations of the glycosylation of total IgG with the level of thyroid autoantibodies or with AITD has been performed.

Our goal was to determine if there are any IgG or PBMC glycan structures associated with the AITD or TPOAb positivity and examine if there are any common heritable factors between AITD and glycan structures. We investigated the association of total serum or plasma IgG glycome composition and PBMC glycosylation in AITD and looked for possible common genetic background in over 3,000 individuals. We found an association between the decreased level of IgG core fucosylation and PBMCs antennary αl, 2 fucosylation with TPOAb level as well as with AITD. We observed the association of significantly affected IgG N-glycan traits with *FUT8* and *IKZF1* genes (responsible for IgG fucosylation), but we could not identify SNPs or a general dysregulation of gene expression in whole blood; suggesting a restricted dysregulation of glycosylation in a subpopulation of B-cells.

## Results

### TPOAb level and AITD are associated with a decreased level of IgG core fucosylation

The presence of autoimmune antibodies is not a definite sign of the AITD, so we wanted to test whether IgG glycosylation status can play a role in active AITD and correlated with the TPOAb level or AITD. We investigated the associations between total plasma IgG glycome composition and peripheral blood TPOAb level and AITD status in 2,297 (988 controls and 1309 TPOAb positive) and 1,191 individuals (988 controls and 203 AITD) respectively from the TwinsUK cohort (Discovery Cohort; **Supplementary Table 1**) using hydrophilic interaction chromatography ultraperformance liquid chromatography (HILIC UPLC). HILIC-UPLC chromatograms were all separated into 24 glycan peaks (GP-Zagreb code, IGP-Edinburgh code, **Fig. 1a**, **Supplementary Table 2**), and the amount of glycans in each peak was expressed as a percentage of the total integrated area. Furthermore, we excluded glycan peak GP3 due to the co-elution with the contaminant and combined GP20 and GP21 to get 22 directly measured N-glycan traits (**Fig. 1c,d; Supplementary Table 2**). As many of these structures share the same features (terminal galactose, terminal sialic acid, core-fucose, bisecting *N*-acetylglucosamine (GlcNAc)), we calculated 53 additional derived traits that average these features across multiple glycans (**Fig. 1c,d; Supplementary Table 2**). Latter structural features were found to be more closely related to individual enzymatic activities in different cellular compartments^29^ and underlying genetic polymorphisms^23^ than the 24 original glycan peaks. As all these structural features partially correlated (**Fig. 1b**), we estimated only 20 independent glycan traits^23^ in our data using the Equation (5) method proposed by Li & Ji (2005)^30^.

**Figure 1.**
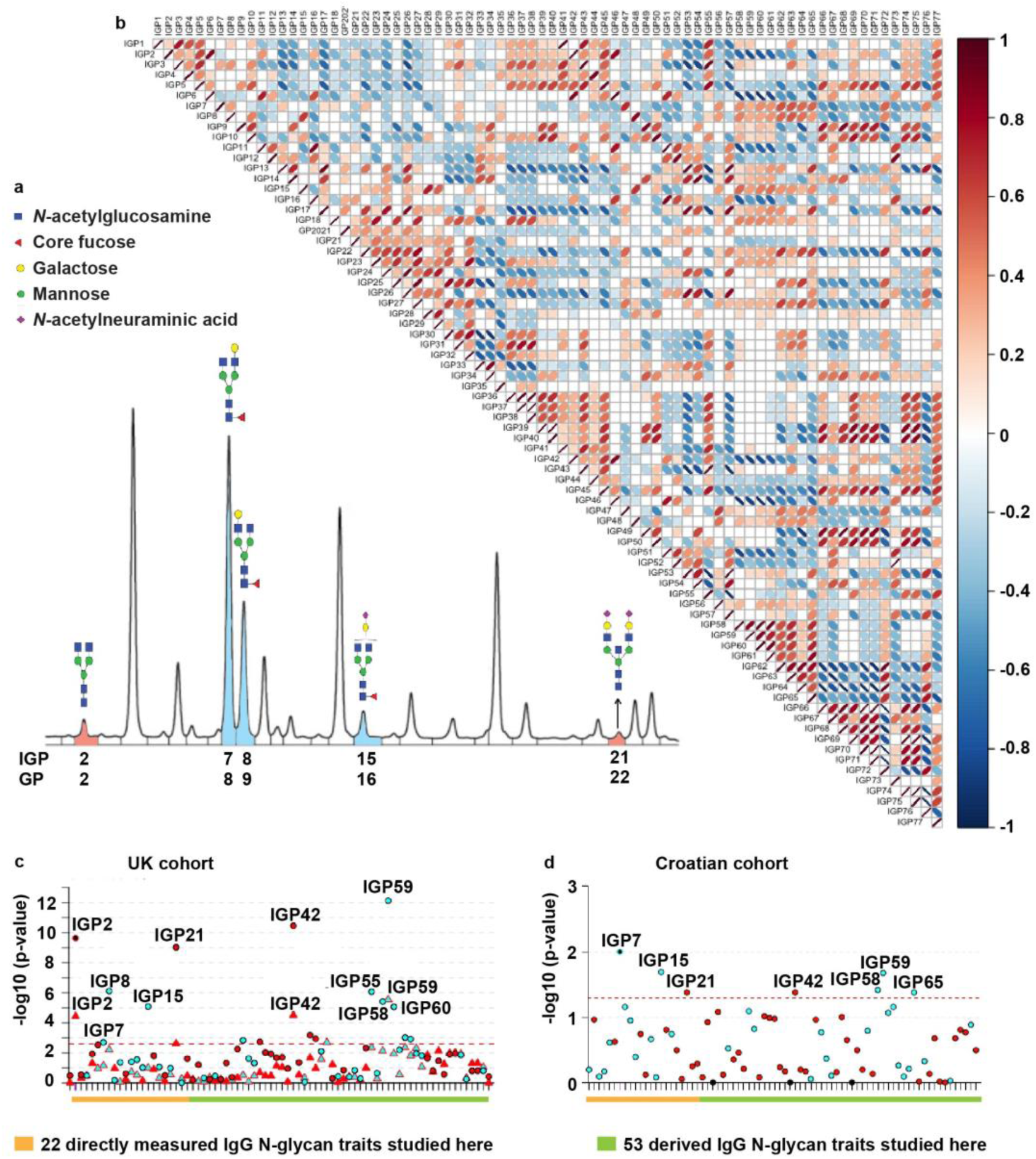
UPLC analysis of the IgG N-glycome and Manhattan plot of IgG N-glycan traits with AITD status and TPOAb level in two cohorts. a) Representative chromatogram showing separation of labeled IgG N-glycan structures into 24 chromatographic IgG glycan peaks (IGPs) is shown with indicated structures of glycans that significantly decreased (blue) or significantly increased (red) with TPOAb level and/or AITD status. b) Correlation matrix between 75 individual IgG N-glycan traits in a general population (the TwinsUK cohort). Image created with R package corrplot. c) Manhattan plot of 75 IgG N-glycan traits (directly measured and derived) in the discovery TwinsUK cohort (Supplementary Table 3 - Sheet 1 and 2). The Manhattan plot is drawn with colors corresponding to the direction of associations (blue=negative, red=positive) and phenotypes are distinguished by the shapes of dots (circle=TPOAb level, triangle =AITD status). The red dashed line corresponds to the level of significance in the discovery cohort (P-value≤2.5×10^−3^). The orange line highlights 22 original glycan peaks detected by HILIC UPLC and analyzed in this study where a chromatogram is represented in Fig. 1a whereas the green line highlights 53 additional IgG N-glycan derived traits. d) Manhattan plot of 75 IgG N-glycan traits for the Croatian replication cohort (TPOAb-positivity; Supplementary Table 3 - Sheet 4). The Manhattan plot is drawn with colors corresponding to the direction of associations (blue=negative, red=positive). The red dashed line corresponds to the level of significance P-value in the replication cohort (0.05). The orange line highlights 22 original glycan peaks detected by HILIC UPLC and analyzed in this study where a chromatogram is represented in Fig. 1a whereas the green line highlights 53 additional IgG N-glycan derived traits. Image created with R package coMET^28^.

The P-value for discovery step (0.05/20= 2.5×10^−3^) was determined using Bonferroni correction for multiple testing with number of independent features^23^. We found eleven directly measured and derived IgG N-glycan traits negatively associated with increased TPOAb level (IGP7, IGP8, IGP15, IGP33, IGP48, IGP56, IGP58, IGP59, IGP60, IGP62 and IGP63) and six directly measured and derived IgG N-glycan traits positively associated (IGP2, IGP21, IGP36, IGP42, IGP45 and IGP46) following Bonferroni correction for multiple testing (P-value <2.5×10^−3^; circle symbol in **Fig. 1c; Supplementary Table 3 - Sheet 1**). Only two derived IgG N-glycan traits from the TPOAb positive group (IGP48 and IGP59) stayed negatively associated with AITD status while three directly measured and derived IgG N-glycan traits (IGP2, IGP21, and IGP42) stayed positively associated with AITD status (triangle symbol in **Fig. 1c; Supplementary Table 3 - Sheet 2**), but no new significant IgG N-glycan traits appeared in the AITD group when compared to the TPOAb positive group of the Twins UK cohort.

Next, we wanted to test if the same TPOAb-associated IgG N-glycan traits are conserved in a cohort of individuals with TPOAb-positivity from Croatia (unknown clinical status). We collected data from 73 control individuals (TPOAb 0.43 (0.53) IU/ml) and 90 case individuals (TPOAb 324.32 (408.46) IU/ml) from Croatia (replication Cohort; Croatian; **Supplementary Table 1**). Four IgG N-glycan traits with negative association (IGP7, IGP15, IGP58 and IGP59) and two IgG N-glycan traits with positive association (IGP21 and IGP42) in the discovery cohort were significantly associated with TPOAb-positivity without established clinical diagnosis in the Croatian cohort (analogous to the ones observed in the TwinsUK discovery cohort) (**Fig. 1d; Supplementary Table 3 – Sheet 4 –** IgG N-glycan traits are bold and blue for negative association or red for positive association).

Investigating the structural composition of directly measured and derived IgG N-glycan traits associated with AITD and the TPOAb level (**Supplementary Table 2**), we observed an increase in IgG N-glycan traits without core fucose (IGP2, IGP21, IGP42, IGP46) and structures with bisecting GlcNAc (IGP36, IGP45) whereas a decrease in IgG N-glycan traits containing core fucose and without bisecting GlcNAc (IGP7, IGP8, IGP15, IGP48, IGP62, IGP63) as well as structures with core fucose regardless of bisecting GlcNAc (IGP58, IGP59, IGP60). Additionally, a decrease in IGP33, which refers to the ratio of all fucosylated (regardless of bisecting GlcNAc) mono- and disialylated structures, was observed. Overall, we found 17 IgG N-glycan traits associated with the TPOAb level, which suggests a decrease in abundance of glycan structures containing core fucose, and five of those structures remain associated with AITD status (**Fig. 1c, d**).

### HT is associated with the decreased level of IgG core fucose and the PBMC antennary α1,2 fucose

Next, we focused our IgG N-glycan trait analysis on individuals with HT. In the TwinsUK Cohort (726 individuals - 675 controls and 51 HT), active HT was identified as TSH>10 mIU/L, or TSH>4 mIU/L accompanied by TPOAb > 100 IU/mL (Discovery cohort; HT; **Supplementary table 1**). To verify the finding, we had an additional cohort of a group of 103 HT patients and 106 control subjects from Poland (Replication cohort; Polish; **Supplementary Table 1)** with clinical manifestation of HT but unknown TPOAb and TSH values at the time of the sampling. Six of the TPOAb associated IgG N-glycan traits remain significant in the UK HT cohort: two IgG N-glycan traits without core fucose (IGP2, IGP42) are increased, while IgG N-glycan traits containing core fucose and without bisecting GlcNAc (IGP8, IGP48), as well as traits representing fucosylation of agalactosylated glycans (IGP59) and percentage of monogalactosylated structures in total neutral IgG N-glycans structures (IGP56) remain decreased (**Fig. 2a**). In contrast, only one of the 17 IgG N-glycan traits that were significant in the discovery cohort (TPOAb level and AITD status; **Supplementary Table 3 – Sheet 3**) was replicated with HT patients in the Polish cohort (IGP48; control versus patients with HT; **Fig. 2b; Supplementary Table 3–Sheet 5**). However, other significant IgG N-glycan traits from the TwinsUK discovery cohort had the same direction for their effect sizes in the Polish replication cohort (**Supplementary Table 3 – Sheet 3, 5;** relevant IgG N-glycan traits are bold). The results from different discovery and replication cohorts suggest a decrease in abundance of glycan structures containing core fucose with the TPOAb level and a reduction of an IgG N-glycan structure with core fucose and galactose in HT.

**Figure 2.**
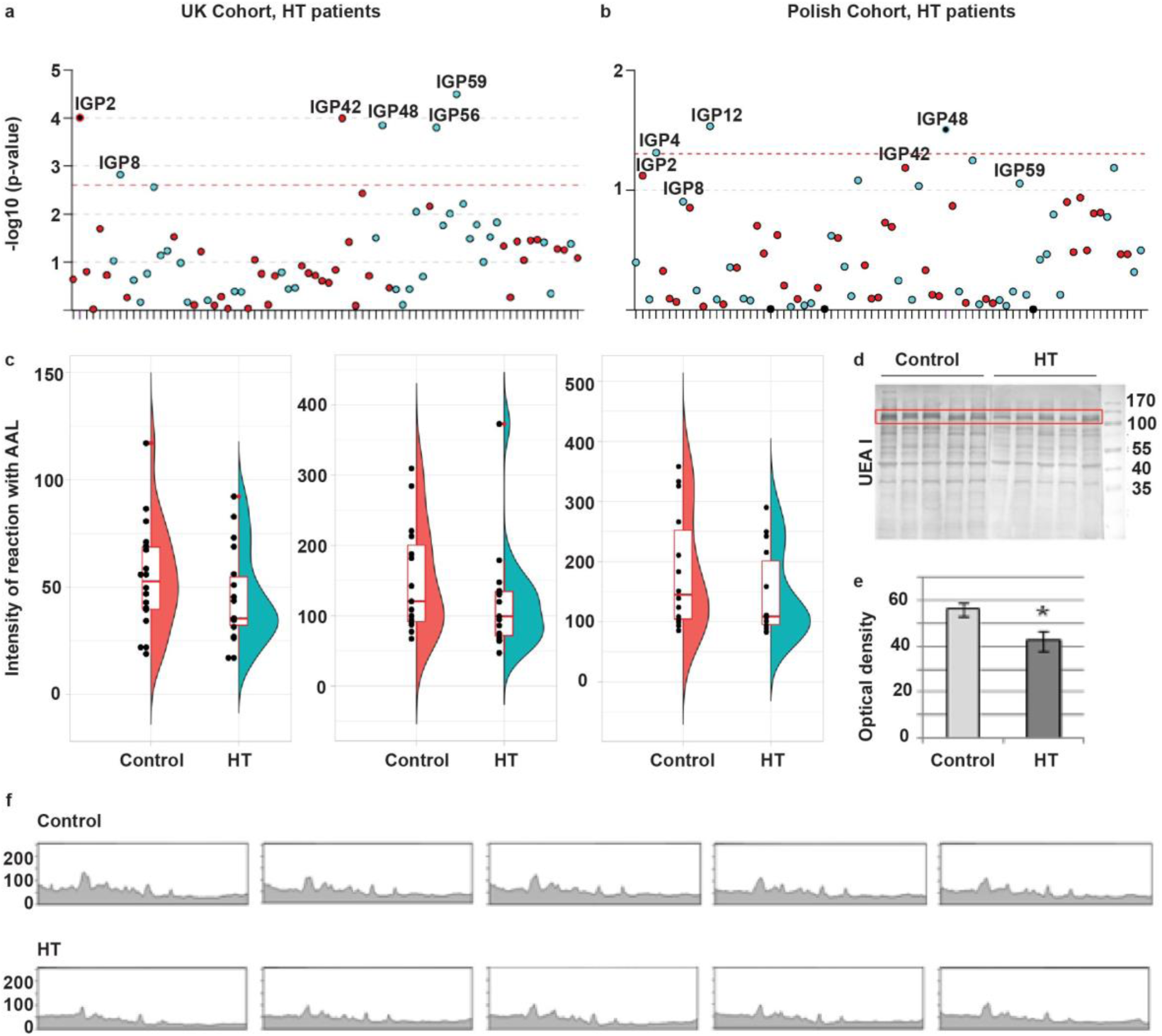
Glycosylation analysis of the human IgG and PBMC proteins from control and HT groups. a) Manhattan plot of 75 IgG N-glycan traits for the HT patients of the UK discovery cohort (**Supplementary Table 3 - Sheet 3**). The Manhattan plot is drawn with colors corresponding to the direction of associations (blue=negative, red=positive). The red dashed line corresponds to the level of significance in the discovery cohort (P-value≤2.5×10^−3^). Image created with R package called coMET. b) Manhattan plot of 75 IgG N-glycan traits for the HT Polish replication cohort (**Supplementary Table 3 - Sheet 5**). The Manhattan plot is drawn with colors corresponding to the direction of association (blue=negative, red=positive). The red dashed line corresponds to the level of significance P-value in the replication cohort (0.05). Image created with R package called coMET. c) Reduction (not significant), of the binding of AAL to IgG core fucose in HT patients compared to control healthy individuals from the Polish cohort in the three batches using lectin blotting assay (P-valueώ0.05; the number of samples per group in each batch is 18,15,14 respectively); **Supplementary Table 4**). The plot combines a flat violin plot, box plot with whiskers and different data. A violin plot is a hybrid of box plot and kernel density plot. Box plot with whiskers represents five summary statistics (the median, the first and third quartiles for two hinges and 1.5 times interquantile range from the hinges for two whiskers). Outliers are labeled by the red dot adjacent to the black dot of the measurement. Image created with R package called ggplot2. d) PBMC protein extracts were resolved on 10% SDS-PAGE gels in reducing conditions. After electrophoresis, the proteins were electrotransferred to a PVDF membrane and stained with the UEA I lectin (n_case_=9 and n_control_=10; **Supplementary Table 5**). The molecular markers are shown on the last lane in kilo Dalton (kDa) (PageRuler Prestained Protein Ladder, 26616, Thermo Scientific). e) Densitometric measurement of the UEA I-positive protein marked in the red frame of Fig. 2d. Duncan’s test was used to compare mean values of optical density of the lectin binding. Statistically significant difference is marked with an asterisk (P-value<0.05). f) The lectin staining of Fig. 2d measured densitometrically.

N-glycosylation of IgG extracted from 20 HT versus 20 healthy donors in Polish cohort (**Supplementary Table 1**) was also examined in three batches by lectin blotting using *Aleuria aurantia* lectin (AAL), specific for α1, 6-linked core fucose. We observed a consistently decreased level of IgG core fucosylation associated with HT status using AAL in the three batches, but none were significant with sample size tested (**Fig. 2c**, P-value> 0.05; **Supplementary Table 4**). We further tested whether the decreased level of core fucosylation in IgG extends to PBMCs and thus whether a specific suite of enzymes associated with a core fucosylation is generally altered in immune cells of AITD patients. N-glycosylation of proteins extracted from PBMC homogenates from HT versus healthy donors (**Supplementary Table 1**) was examined by lectin blotting using seven lectins with different sugar specificity: AAL, *Galanthus nivalis* lectin (GNL), *Mackia amurensis* lectin (MAL-II), *Phaseolus vulgaris* erythroagglutinin (PHA-E), *Phaseolus vulgaris* leucoagglutinin (PHA-L), *Sambucus nigra* agglutinin (SNA), and *Ulex europaeus* agglutinin (UEA I). Lectin blotting with UEA l identified a significant reduction of antennary αl,2 fucose-in patients with HT (**Fig. 2d**) whereas blotting with AAL, which preferentially binds core fucose (α1,6), as well as all other lectin blotting assays, showed no difference in the content of particular glycan species between HT and control group (**Supplementary Fig. 1**, **Supplementary Table 5**). Coomassie Brilliant Blue staining control of total protein amount confirmed that there was no significant difference in protein profiles between control and HT samples (**Supplementary Fig. 2**). To summarize, we found six significant glycans in the UK HT discovery cohort that were also associated with the TPOAb level (one was replicated in the Polish cohort) and a decrease of PBMC antennary α1,2 fucose associated with HT.

### AITD and IgG N-glycan traits are highly heritable, but no shared genetic effect between them could be determined

We and others identified several loci associated with IgG and plasma glycosylation and autoimmune conditions in the previous GWAS^24,25,31,32^, so we wanted to relate those findings to the genetic background of the subjects in this study to determine whether there is a shared heritability of AITD and IgG N-glycan traits. We estimated the proportions of genetic and environmental variance in the phenotypic variation of AITD status and different IgG N-glycan traits using the twin data from the TwinsUK cohort and the maximum likelihood variance component model fitting, also known as a structural equation-modeling or ADCE model^33^. The best models of AITD status and TPOAb-positivity estimated were the DE model - with only a dominance genetic variance (D) and a unique environmental variance (E) - with about 57-63% of heritability (**Supplementary Table 6**), as previously estimated in a Danish twin cohort^25-27^. However, since the TPOAb does not follow a normal distribution, we could not predict with accuracy the best ADCE model. Regarding IgG N-glycan traits, most of IgG N-glycan traits have AE (model with only an additive genetic variance (A) and a unique environmental variance (E)) for the best model and the average heritability for all IgG N-glycan traits was of 55.15% (CI; min=50.54; max=59.74; **Supplementary Table 2**). Given that the heritability for AITD and TPOAb-positivity predicts only a dominance genetic variance as a fraction of genetic variance, and the heritability of IgG N-glycan traits was estimated to be composed with mostly additive genetic variance, the portion of shared genetic variances between AITD and IgG N-glycan traits using a bivariate heritability analysis^34^ could not be determined.

### Lead SNPs associated with IgG glycan FA2G1S1 and TPOAb-positivity fall in the same high LD nearby HCP5 gene

Next, we investigated whether specific genetic variants determined by previous GWASs could explain some genetic components potentially shared between AITD status and IgG N-glycan traits. We compared the results from internal GWASs of IgG N-glycan traits in the TwinsUK cohort (around 4,500 individuals)^24^ and previous GWASs of IgG N-glycan traits^23^ with results from the GWAS catalog for thyroid diseases and functions including GD, TPOAb-positivity, hypothyroidism, hyperthyroidism, levels of thyroid hormones^37^ (**Supplementary Table 7**). No lead SNPs could be found to be associated with at least one of thyroid phenotype and one of IgG N-glycan traits. By exploiting linkage disequilibrium (LD; r^2^>0.8) around the lead SNPs detected by these previous GWASs, we observed that the lead SNP of IGP15 (rs3094014) fell in the same high LD block around *HCP5* (HLA Complex P5) gene on the chromosome 6 within the MHC class I region, as the lead SNP of TPOAb-positivity (rs3094228) (**Fig. 3**). Moreover, we found that the lead SNP of IGP14 and IGP54 (rs199442) clustered with hypothyroidism SNP (rs77819282) in the same high LD around the *NSF* (N-Ethylmaleimide Sensitive Factor, Vesicle Fusing ATPase) gene on the chromosome 17. Neither IGP14 nor IGP54 was associated with AITD and TPOAb-positivity. However, one of the significant IgG N-glycan traits, IGP15 also known as IgG glycan FA2G1S1, is correlated with the TPOAb level in our current study. Both SNPs relevant to both IGP15 and TPOAb-positivity (rs3094014 and rs3094228), as well as the other SNPs in this high LD (rs3099840, rs3131620, rs3128987), have a cell-type specific expression quantitative trait loci (eQTLs). For example, rs3094014 is an eQTL for *C4A* (thyroid, P-value=2.2×10^−14^; whole blood P-value=1.5×10^−7^), *CYP21A1P* (thyroid, P-value=4.7×10^−12^; whole blood P-value=2.0×10^−9^), *HLA-C* (thyroid, P-value=1.9×10-6), *HCP5* (thyroid, P-value=1.1×10^−9^; whole blood, p=1.2×10^−9^), and *CYP21A2* (whole blood, P-value=7.1×10^−8^), whereas rs3094228 is associated with expression of *C4A* (thyroid, P-value=3.4×10^−13^; whole blood P-value=1.07×10^−7^), *CYP21A1P* (thyroid, P-value=1.1×10^−9^; whole blood P-value=3.79’10^−8^), *HLA* (PBMC, P-value= 2.09’10^−7^ to 8.77’10^−23^), *HCP5* (thyroid, P-value=1.3’10^−9^; whole blood, P-value=1.86×10^−8^), and *CYP21A2* (whole blood, P-value=3.24×10^−7^) ^38-40^. This data indicates that SNPs associated with both IGP15 and TPOAb-positivity clustered around *HCP5* gene alter the expression of genes in the thyroid cells and whole blood cells and fall in DNase sensitive site in T-cells and B-cells.

**Figure 3.**
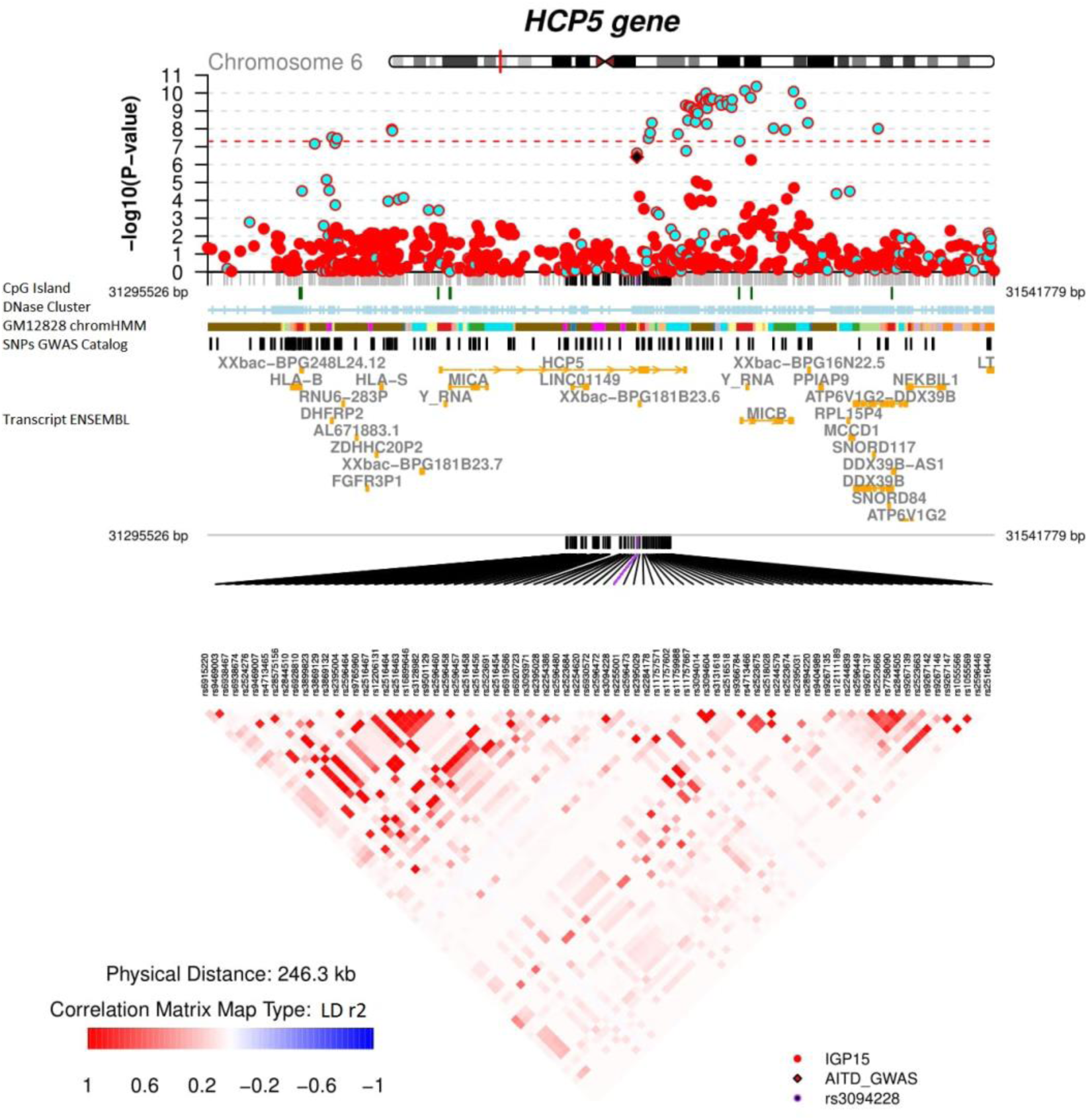
coMET plot for SNP association data on IGP15 in whole blood and TPOAb-positivity in the *HCP5* gene region. Regional Manhattan plot from a GWAS of *IGP15* performed on the TwinsUK cohort and TPOAb-positivity from meta-analysis GWAS^35^. The regional Manhattan plot is drawn with colors corresponding to the direction of association (blue=negative, red=positive), the solid diamond symbol (black) represents the SNP rs3094228, which was associated with TPOAb-positivity^35^. The red dashed line corresponds to the level of significance (P-value=5×10^−8^). Annotation tracks corresponding to CpG island (UCSC), Encode Dnase cluster (UCSC), GM12878 Chromatin State Segmentation by HMM from Encode/Broad (UCSC), SNPs from GWAS Catalog (UCSC) and transcripts from ENSEMBL are displayed below the regional Manhattan plot. An LD matrix computed from genotypes of twins from the TwinsUK cohort (r^2^ using PLINK software^36^) is also provided. For Encode/Broad ChromHMM the colors correspond to: Red = 1. Active Promoter, Pink = 2. Weak Promoter, Purple = 3. Poised Promoter, Orange = 4 Strong Enhancer, Lilac = 5. Strong Enhancer, Yellow = 6. Weak Enhancer, Light Orange = 7. Weak Enhancer, Blue=8.Insulator, Light Green = 9. Txn Transition, Green = 10. Txn Elongation, Cyan = 11. Weak Txn, Magenta = 12. Repressed, Brown = 13. Heterochrom/lo. Image created with R package called coMET^28^.

### Enrichment of IgG N-glycan traits associated with two enzymes, Fut8 and Ikzf1, essential for the core fucosylation of IgGs in TPOAb positive subjects

We then focused on the nine groups of genes that were found to be associated with modifications in IgG glycosylation pattern from the previous GWASs^23^, and we evaluated which groups of genes associated with IgG glycosylation patterns that are over-represented among IgG N-glycan traits associated with TPOAb positivity and AITD in the current study. Using the 17 significant IgG N-glycan traits in the discovery cohort and out of the nine groups of genes, we observed an enrichment of IgG N-glycan traits associated with *FUT8* and *IKZF1* genes (P_*FUT8*_=3.52×10^−3^; odd ratio_*FUT8*_=7.28 [IGP2, IGP42, IGP46, IGP58, IGP60, IGP63]; P_*IKZF1*_=8.74×10^−4^; odd ratio_*IKZF1*_=9.19 [IGP2, IGP42, IGP46, IGP58, IGP59, IGP60, IGP62], **Fig. 4**). Both were associated with core fucosylation of IgGs regardless of bisecting GlcNAc^23^. When we repeated the analysis with only the subset 7 replicated IgG N-glycan traits replicated in Croatian cohort, the enrichments became non-significant (P_*FUT8*_=7.41×10^−2^; odd ratio_*FUT8*_=4.85 [IGP42, IGP58]; P_*IKZF1*_=9.21×10^−2^; odd _ratio*IKZF1*_=4.30 [IGP42, IGP58, IGP59] respectively). On the other hand, none of the 17 significant IgG N-glycan traits, and so of 7 replicated IgG N-glycan traits, was associated with the *LAMB1* gene which was associated with core fucosylation of IgGs with the bisecting GlcNAc (P_*LAMB1*_=1; odd ratio_*LAMB1*_=0) in Lauc et al., 2013^23^. Overall, we observed an enrichment of IgG N-glycan traits associated with two genes, *FUT8* and *IKZF1*, essential for the formation of IgG core fucose in TPOAb positive subjects.

**Figure 4.**
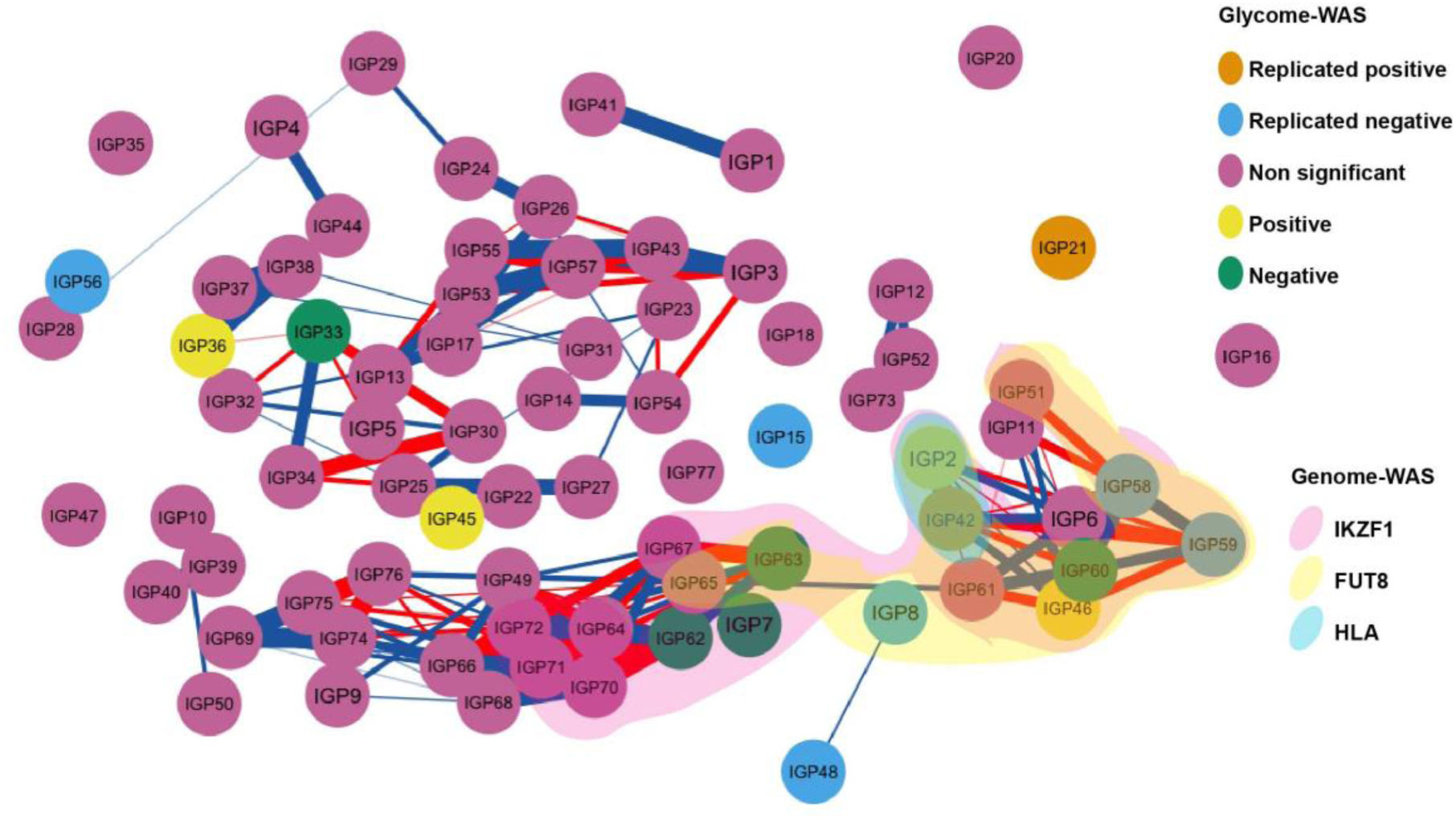
Enrichment in AITD patients of IgG N-glycan traits associated with *IKZF1* and *FUT8* genes. Correlation network between 75 IgG N-glycan traits, highlighting 17 IgG N-glycan traits associated with AITD and TPOAb level and those associated with SNPs close to *HLA*, *FUT8*, *IKZF1* genes. Nodes represent different IgG N-glycan traits (the color of nodes is related to the associations performed in discovery and replicated cohorts) and connections show the Pearson correlation between IgG N-glycan traits (only correlations more than 0.75 are visualized, red for negative correlation and blue for positive correlation, the strength of correlation is represented by the width of edge). The three clouds above of nodes cluster the different IgG N-glycan traits previously found to be associated with SNPs within/close to *HLA* gene (blue cloud), the *FUT8* gene (yellow cloud) and *IKZF1* gene (red cloud)^23^. Image created with R package qgraph.

### The decreased level of IgG core fucose and PBMC antennary α1,2 fucose in AITD is not associated with a modification of gene expressions in circulating whole blood cells

A decrease in IgG core fucose, PBMC antennary α1,2 fucose, and no lead SNPs in the fucosylation genes associated with TPOAb positivity or AITD could all be explained by general dysregulation in gene expression belonging to fucosylation pathways. To explore this possibility, we performed transcriptome-wide association scan (TWAS) of the TPOAb level and AITD status on up to 199 individuals from the TwinsUK cohort^41^. We further performed TWAS of 75 N-glycan traits on general TwinsUK cohort regardless of TPOAb level or AITD status (326 individuals) (**Supplementary Table 1 - sheet 2**). No significant modification of gene expressions (P-value <5.82×10-7) was genome-wide associated with AITD and TPOAb at the exon level, even for subsets of genes such as those belonging to the pathways of fucosylation, including *FUT1*, *FUT2*, *FUT8*, *LAMB1*, and *IKZF1*. The decreased level of IgG and PBMC fucosylation in AITD could not be confirmed to be a consequence of a general depletion of fucosylation pathways in whole blood cells. However, in general population regardless for AITD status or TPOAb level (**Supplementary Table 8**), IGP58 and IGP60 were associated with ENSG00000211949.2, the main exon of *IGHV3*-*23*, immunoglobulin heavy variable 3-23 present on chromosome 14 (beta=0.25; P-value<4.9×10^−7^ for both IgG) while IGP45 was associated with ENSG00000101986.7, the 7^th^ exon of *ABCD1* gene (beta=-0.35; P-value=2.43×10^−7^) and with ENSG00000072310.10, the 10th exon of *SREBF1* gene (beta=-0.34; P-value=5.18×10^−7^). Decreased core fucosylation and PBMC antennary α1,2 fucose are not accompanied by the alteration of gene expression in PBMC.

## Discussion

The role of the autoantibodies in the development of AITD is still unknown. AITD are among the most frequent autoimmune disorders occurring in almost 5% of general population^1^. If individuals harboring positive antibodies against thyroid proteins but without active disease are included, that adds up to 15% of the general population affected by thyroid autoimmunity^4^. Three central thyroid autoantibodies (TPOAb, TgAb, TSAb) were identified in patients with AITD, and their role in AITD is still unclear. Since the presence of TPOAb does not necessarily indicate active AITD, but IgG glycosylation was previously shown to be an essential factor in regulating autoantibody function in autoimmune diseases^18,19^, we examined the IgG glycosylation status in AITD patients and TPOAb positive individuals. This study is the first that investigated the potential association of IgG and PBMC glycosylation with AITD and TPOAb level, genetic background between them and hypothesized about the causality of these relationships.

We observed a decrease in IgG core fucose level, which presented as altered levels of 17 IgG N-glycan traits in the whole blood of individuals with TPOAb, and depletion of PBMC antennary α1,2 fucose in patients with HT. Six out of seventeen significant IgG N-glycan traits for TPOAb positivity were replicated in a Croatian cohort. Only one out of these six IgG N-glycan traits was replicated in a Polish cohort, however, the other five significant structures from the UK cohort showed the same trend in the Polish cohort. Several healthy individuals in the Polish cohort have high thyroid autoantibodies (TgAb or TPOAb), but lower than the threshold set by the manufacturer for reporting of clinically elevated levels, whereas some HT patients have low thyroid autoantibodies. Additionally, all HT patients are under treatment with thyroid drugs that could affect their levels of thyroid autoantibodies^42,43^. Therefore, a possible reason why only one significant glycan feature was replicated in the Polish HT cohort might be the level of TPOAb. Unfortunately, there are not enough samples to stratify the Polish cohort according to TPOAb level or to classify the Croatian cohort according to clinical diagnosis to test the importance of TPOAb level in findings using UPLC data. However, when we reduced the case group from Croatia to 83 individuals with very high TPOAb (>100 IU/ml in Roche assay, the same criteria to define AITD status in the TwinsUK cohort), the associations for the six replicated IgG N-glycan traits became more significant while the beta values remained similar to the preceding analysis (**Supplementary Table 3 – Sheet 4**). Using lectin blotting in the Polish cohort, we validated the reduction of core-fucosylated IgG in HT patients, but due to the lack of data on TPOAb levels for HT patients (i.e. analyzed in the same blood collected for IgG glycosylation analysis), we could not confirm the association with TPOAb level rather than AITD status. Overall, our results suggest that the decreased fraction of core-fucosylated IgG could be more related to the level of TPOAb than the status of AITD.

Several studies showed that 95 *%* of IgG N-glycan structures in a healthy individual have core fucose and it acts as a “safety switch”, attenuating potentially harmful ADCC^44-48^ Using *FUT8* knockout Chinese hamsters, one previous study produced de-fucosylated CD20 antibodies and showed that afucosylated antibodies have a much higher affinity for FcγRIIIa (CD16a) - an immunoglobulin receptor distributed on natural killer (NK) cells, macrophages, and γδ T cells. They enhanced ADCC over 100-fold more than fucosylated CD20 antibodies^49^. Production of antibodies without the core fucose has recently revolutionized antibody therapies, by providing substantially enhanced ADCC^50,51^. Thus, the deficiency of IgG core fucose observed in the people with AITD and TPOAb-positivity in our data and the activities of IgG core fucose on the immune system previously identified, suggest an increase of ADCC in TPOAb positive individuals.

Interestingly, ADCC was previously reported in AITD without restriction to subgroups of patients, however, more patients with HT than with GD presented ADCC activities^52,53^. Although no strong correlation of TPOAb level (total IgG or IgG subclasses) measured by enzyme-linked immunosorbent assay (ELISA) with ADCC activity could be found in previous studies, and other thyroid antigens besides TPO could be involved in ADCC, TPO was described as the primary antigen involved in the thyroid ADCC^52-55^. Deglycosylation of TPOAb was shown to reduce the binding to FCγRS and thus inhibit the apoptosis of thyroid cancer cells via complement-dependent cytotoxicity (CDC) and cytotoxicity via ADCC^54^. Although IgG glycosylation studied in the present paper is of total and not antigen-specific IgG, we found a relationship between TPOAb level and IgG core fucose, and we suggest that the depletion of core fucose observed in TPOAb circulating in blood could enhance their cytotoxicity activities on thyrocytes through ADCC (**Fig. 5**).

**Figure 5.**
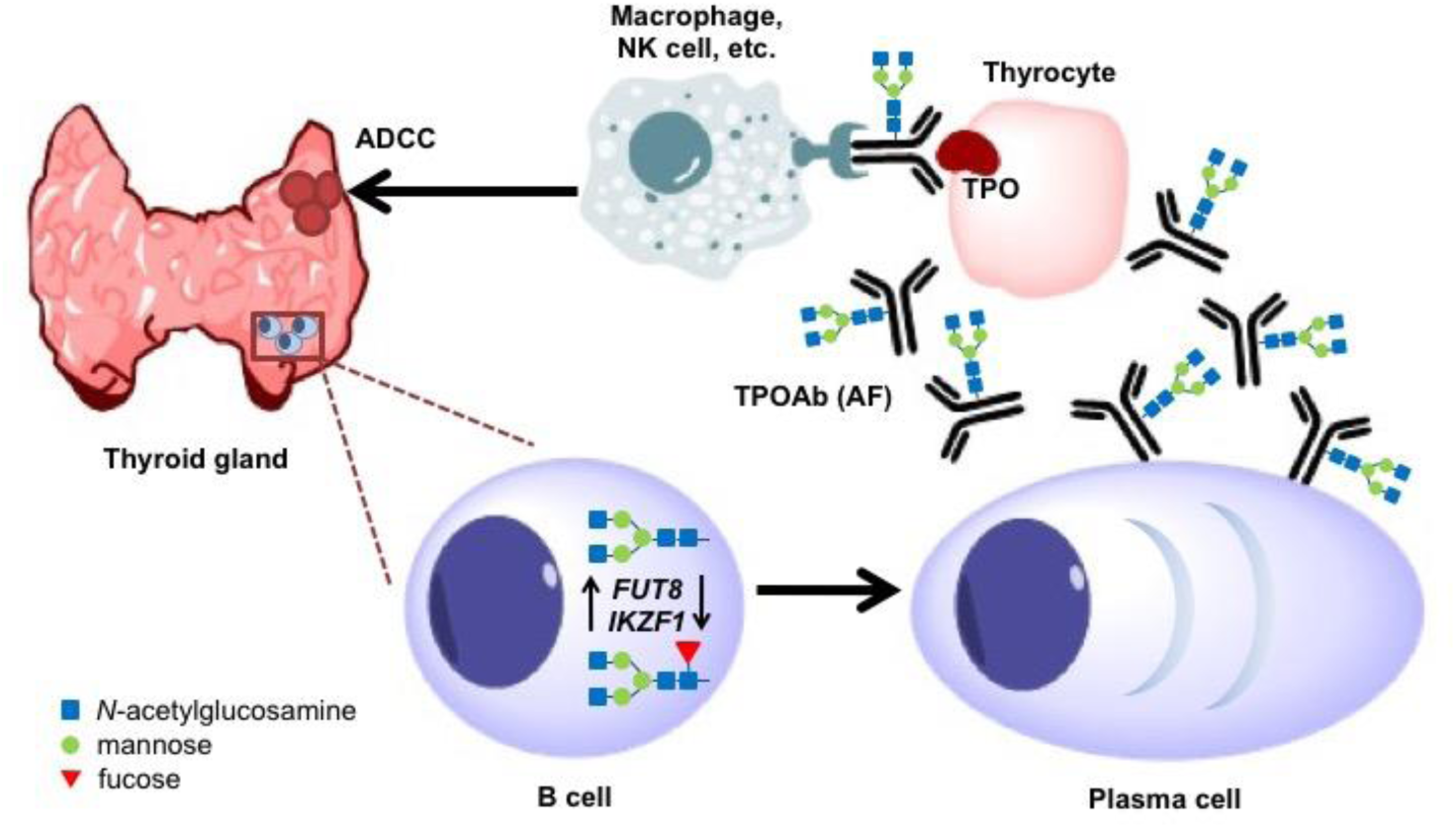
Hypothetic model of the possible role of the glycosylation of TPOAb driven by tissue-specific *FUT8* and *IKZF1* expression in AITD patients. Due to the lack of the significant deregulation of the expression of *FUT8* and *IKZF1* genes in whole blood of AITD patients, these genes could have an aberrant expression in a tissue-specific manner. It was previously suggested that TPOAb producing B-cells are generated in the germinal centers that are appearing in the thyroid gland of the AITD patients. Those cells could have aberrant expression of *FUT8* and *IKZF1* genes resulting in afucosylated (AF) TPOAb antibodies that are stimulating potentially harmful ADCC directed against the thyroid epithelial cells (thyrocytes), and therefore contributing to the AITD development.

Furthermore, taking advantage of the twin study design of the discovery cohort, we tested shared genetic and environmental effects between different significant glycan features found to be associated with TPOAb positivity and AITD in this study. Surprisingly, although previous GWASs on IgG N-glycan traits showed several SNPs also associated with autoimmune diseases^23,56^, no shared genetic variance or lead SNPs could be detected between IgG N-glycan traits with AITD status or the TPOAb level. In exploring in LD of different lead SNPs associated with thyroid diseases and IgG N-glycan traits, we found that a SNP associated with IGP14 and IGP54 (rs199442) clustered with hypothyroidism SNP (rs77819282) in the same high LD block around NSF, but since neither IGP14 or IGP54 were found to be associated with TPOAb and AITD, *NSF* could potentially have a pleiotropic effect on hypothyroidism that we could not address with current data. SNPs rs3094014 and rs3094228 appear to be associated with IGP15 and AITD respectively, are in high LD, are located in the *HCP5* gene region and indeed are eQTLs for *HCP5* and other genes associated with immune response. These two lead SNPs and four other of the same LD block fall in a regulatory element, more precisely an enhancer and an open-chromatin in B-cell and thyroid tissue. Even though we identified two SNPs significantly associated with the IgG N-glycan structure IGP15, the cross-sectional nature of the cohorts and lack of independence of previous GWASs with the TwinsUK cohort do not allow conclusions to be drawn on the causal relationship between IgG core fucosylation and AITD.

Based on the findings of GWASs performed on the IgG N-glycan traits^23^, we also showed an enrichment of the afucosylated IgG N-glycan traits that are associated with *FUT8* and *IKZF1* genes. Both genes were considered leading players in the core fucosylation of IgGs, although the mechanism of *IKZF1* gene in the fucosylation is still unclear^23^. The *IKZF1* gene encodes a transcription factor belonging to the family of zinc-finger DNA-binding proteins associated with chromatin remodeling and regulating lymphocyte differentiation. Interestingly, several SNPs around the *IKZF1* gene have been associated with autoimmune diseases based on the GWASs’ findings; including systemic lupus erythematosus^57^, Crohn’s disease^58^, inflammatory bowel disease^58^, and other diseases such as acute lymphoblastic leukemia^59^. On the other hand, the *FUT8* gene encodes fucosyltransferase 8, a known enzyme catalyzing the addition of fucose in αl, 6 linkage to the first GlcNAc residue (core fucose), but no SNPs around *FUT8* genes have been associated with immune phenotypes other than IgG and plasma N-glycan structures from the previous GWASs^23,31,32,37^. However, even though these two fucosylation enzymes were found to be associated with the significant N-glycan structures from the present study, we could not find a direct association of the SNPs falling within or near their genomic regions with AITD or TPOAb at the current sample size. Since FUT8 and IKZF1 are essential players in the regulation of the core fucose expression and as we found no identifiable genomic mutations occurring in or near those genes in TPOAb positive individuals or AITD patients, we went on to examine the expression of those genes in whole blood.

The transcriptomic analysis in the whole blood cells performed in the TwinsUK cohort suggested that the general decrease of fucosylation associated with AITD and TPOAb level was not a consequence of dysregulated gene expression of known fucosylation genes in the circulating blood. Furthermore, with the current data, we were also not able to find a dysregulation in PBMC transcriptome associated with AITD that is linked to PBMC fucosylation. We are unable to hypothesize about the functionalities and pathways of production of PBMC antennary α1,2 fucose as up to now, PBMC glycosylation is not well characterized due to the difficulty in the isolation of a sufficient amounts of individual types of PBMC (e.g. B-cells and CD4+ T-cells) for glycan analysis. In comparison, IgG glycan structures are well studied as the IgG comprises approximately 75% of total serum immunoglobulins. Considering we measured IgG glycosylation from whole blood and found the enrichment of afucosylated N-glycan traits, the statement that we found no general dysregulation of the fucosylation pathway in whole blood might seem confusing at first. However, although antibodies circulate and are detected in the blood, most plasma cells that produce circulating IgGs are located in germinal centers of secondary lymphoid organs like the spleen and lymph nodes^60^. In the case of AITD, the production of antibodies against thyroid components was suggested to happen directly in the thyroid gland where formations of germinal centers have occurred in patients with AITD^61-63^. Consequently, in addition to the previous detection of germinal centres in the thyroid gland of patients with AITD, our finding from transcriptomic analysis suggests that the IgG glycosylation pattern observed in circulating blood of people with AITD status is not directly associated with immune cells in whole blood, but could be a dysregulation of specific immune cell-types such as thyroid-derived lymphocytes that produce thyroid autoantibodies^64^ (**Fig. 5**).

In this study, we used fixed volumes of total IgG solution and not the same initial mass of IgG for the UPLC analysis of released glycans. Additionally, IgG glycosylation was analyzed on the level of total IgG, not on the level of a specific antibody. This approach was chosen because of the fixed amount of residual salts and the difficulty in obtaining enough material for analysis of IgG glycosylation of a particular antibody (e.g. absence or low concentration of thyroid autoantibodies in the total IgG in healthy individuals). Preparations of reference of TPOAb level were made from a pool of serum from patients with AITD and were prepared and lyophilized 35 years ago and use the international unit per milliliter (IU/mL) as the reference unit. Unfortunately, there is no simple way to convert TPOAb level from IU/mL provided by the assays in the present study to a concentration expressed as microgram per milliliter (μg/ml) and check if the concentration of total IgG is altered by the secretion of thyroid autoantibodies. The concentrations of TPOAb and TgAb were previously measured (up to 1.4 mg/ml for TPOAb and 0.7 mg/ml for TgAb) ^65-67^. Knowing that normal IgG concentration range is 5.1-15.8 mg/ml, in extreme cases, thyroid autoantibodies could represent up to 50% of total IgGs, and their presence in the blood can also increase the serum levels of total IgGs^68,69^. Thus, the secretion of thyroid antibodies can alter the concentration of total IgG in AITD.

If the modification of IgG glycosylation observed in the current study is the consequence of increased TPOAb concentration in total IgGs, the hypothesis that the depletion of IgG core fucose is a biomarker of TPOAb and its activity, and thereby a risk factor of harmful ADCC in the thyroid cells triggered by TPOAb (**Fig. 5**), becomes more likely. The cross-sectional nature of cohorts in this study and lack of independence of previous GWASs with the TwinsUK cohort do not allow conclusions on the causal relationship between IgG core fucosylation and AITD. Therefore, further analysis needs to be performed to describe genes and mechanisms playing in IgG and PBMC glycosylation in AITD in more specific tissues and cell-types to test this hypothesis. Studies looking at the expression of the known fucosylation genes in immune cells from thyroid tissue of healthy subjects, TPOAb positive individuals, and AITD patients might be especially valuable in dissecting the role of the fucosylation in the AITD disease genesis. However, to determine the causality between TPOAb level, fucosylation, ADCC and possibly other factors involved in the AITD genesis, prospective longitudinal studies that look at these traits before and after the diagnostic of AITD, as well as Mendelian randomization^70,71^ in large independent cohorts would be required.

## Conclusion

This study identified for the first time an association of decreased level of IgG core-fucosylation and PBMC antennary α1,2 fucosylation with TPOAb level and AITD. This is also the first time that a decrease of fucosylation on IgG and PBMC is associated with one autoimmune disease and one of its biomarkers. Our findings could not be explained by common genetic background or general dysregulation of gene expression in the whole blood. However, drawing from the knowledge generated in a large number of previous studies, we could speculate that this reduction of IgG core fucose could be a consequence of tissue-specific aberrant gene expression. Moreover, we hypothesized that the decreased IgG core fucosylation could be a novel risk factor for potentially harmful ADCC in the thyroid gland, associated with TPOAb and AITD. Further studies of the glycosylation of thyroid autoantibodies and their interactions with other immune features (e.g., immune cell-types, secreted proteins) and thyroid cells may be helpful to elucidate the potential role of autoantibodies and their glycosylation patterns in the pathogenesis and the treatment of thyroid diseases.

## Acknowledgment

- Massimo Mangino, Jonas Zierer, and Maxim Freydin, King’s College London, UK, for discussion in the analysis of glycosylation data.
- TwinsUK cohort. The study was funded by the Wellcome Trust; European Community’s Seventh Framework Programme (FP7/2007-2013). The study also receives support from the National Institute for Health Research (NIHR)-funded BioResource, Clinical Research Facility and Biomedical Research Centre based at Guy’s and St Thomas’ NHS Foundation Trust in partnership with King’s College London and the Australian National Health and Medical Research Council (PG 1087407).
- Polish cohort: The study was supported by the grants from the Jagiellonian University (PBMC glycosylation, K/DSC/001749; IgG glycosylation, K/DSC/002341).
- IgG glycan analysis in the three cohorts was performed in Genos Glycoscience Research Laboratory and partly supported by the European Union’s Horizon 2020 grants SYSCID (grant agreement No. 733100), IMforFUTURE (grant agreement No. 721815), GlySign (grant agreement No. 722095) and by the European Structural and Investment Funds IRI grant (#KK.01.2.1.01.0003) and Croatian National Centre of Research Excellence in Personalized Healthcare grant (#KK.01.1.1.01.0010).

## References

1. Wang, B., Shao, X., Song, R., Xu, D. & Zhang, J.-A. The Emerging Role of Epigenetics in Autoimmune Thyroid Diseases. Front. Immunol. 8, 174–13 (2017).

2. De Groot, L. Graves’ Disease and the Manifestations of Thyrotoxicosis. Thyroid Disease Manager 1–77 (2015).

3. Mariotti, S., Caturegli, P., Piccolo, P., Barbesino, G. & Pinchera, A. Antithyroid Peroxidase Autoantibodies in Thyroid Diseases. Journal of Clinical Endocrinology and Metabolism 71, 661–669 (1990).

4. Hollowell, J. G. et al. Serum TSH, T sub4/sub, and Thyroid Antibodies in the United States Population (1988 to 1994): National Health and Nutrition Examination Survey (NHANES III). The Journal of Clinical Endocrinology & Metabolism 87, 489–499 (2002).

5. Seeling, M., Brückner, C. & Nimmerjahn, F. Differential antibody glycosylation in autoimmunity: sweet biomarker or modulator of disease activity? Nature Reviews Rheumatology 13, 621–630 (2017).

6. Kobata, A. The N-linked sugar chains of human immunoglobulin G: their unique pattern, and their functional roles. Biochimica et biophysica acta 1780, 472–478 (2008).

7. Shade, K.-T. & Anthony, R. Antibody Glycosylation and Inflammation. Antibodies 2, 392–414 (2013).

8. Novokmet, M. et al. Changes in IgG and total plasma protein glycomes in acute systemic inflammation. Scientific Reports 4, 4347 (2014).

9. Marth, J. D. & Grewal, P. K. Mammalian glycosylation in immunity. Nature Rev Immunology 8, 874–887 (2008).

10. Maverakis, E. et al. Glycans in the immune system and The Altered Glycan Theory of Autoimmunity: a critical review. J. Autoimmun. 57, 1–13 (2015).

11. Subedi, G. P. & Barb, A. W. The immunoglobulin G1 N-glycan composition affects binding to each low affinity Fc γ receptor. MAbs 8, 1512–1524 (2016).

12. Arnold, J. N., Wormald, M. R., Sim, R. B., Rudd, P. M. & Dwek, R. A. The impact of glycosylation on the biological function and structure of human immunoglobulins. Annu. Rev. Immunol. 25, 21–50 (2007).

13. Bondt, A. et al. Immunoglobulin G (IgG) Fab glycosylation analysis using a new mass spectrometric high-throughput profiling method reveals pregnancy-associated changes. Molecular & cellular proteomics: MCP 13, 3029–3039 (2014).

14. Spiegelberg, H. L., Abel, C. A., Grey, H. M. & Fishkin, B. G. Localization of the Carbohydrate within the Variable Region of Light and Heavy Chains of Human γG Myeloma Proteins. Biochemistry 9, 4217–4223 (1970).

15. Abel, C. A., Spiegelberg, H. L. & Grey, H. M. Carbohydrate content of fragments and polypeptide chains of human .gamma.G-myeloma proteins of different heavy-chain subclasses. Biochemistry 7, 1271–1278 (1968).

16. van de Bovenkamp, F. S., Hafkenscheid, L., Rispens, T. & Rombouts, Y. The Emerging Importance of IgG Fab Glycosylation in Immunity. The Journal of Immunology 196, 1435–1441 (2016).

17. Jennewein, M. F. & Alter, G. The Immunoregulatory Roles of Antibody Glycosylation. Trends in Immunology 38, 358–372 (2017).

18. Vučkovic, F. et al. Association of systemic lupus erythematosus with decreased immunosuppressive potential of the IgG glycome. Arthritis and Rheumatology 67, 2978–2989 (2015).

19. Trbojević-Akmačić, I. et al. Inflammatory Bowel Disease Associates with Proinflammatory Potential of the Immunoglobulin G Glycome. Inflammatory Bowel Diseases 21, 1 (2015).

20. Rombouts, Y. et al. Glycosylation of immunoglobulin G determines osteoclast differentiation and bone loss. Nature Communications 6, 6651 (2015).

21. Yuan, S. et al. Changes in anti-thyroglobulin IgG glycosylation patterns in Hashimoto’s thyroiditis patients. The Journal of clinical endocrinology and metabolism 100, 717–724 (2015).

22. Zhao, L. et al. Glycosylation of sera thyroglobulin antibody in patients with thyroid diseases. European Journal of Endocrinology 168, 585–592 (2013).

23. Lauc, G. et al. Loci associated with N-glycosylation of human immunoglobulin G show pleiotropy with autoimmune diseases and haematological cancers. PLoS Genet 9, e1003225 (2013).

24. Shen, X. et al. Multivariate discovery and replication of five novel loci associated with Immunoglobulin G N-glycosylation. Nature Communications 1–10 (2017). doi:10.1038/s41467-017-00453-3

25. Brix, T. H., Kyvik, K. O. & Hegedüs, L. A Population-Based of chronic Autoimmune Hypothyroidism in Danish Twins. Jcem 85, 536–539 (2000).

26. Brix, T. H., Kyvik, K. O., Christensen, K. & Hegedüs, L. Evidence for a major role of heredity in Graves’ disease: a population-based study of two Danish twin cohorts. The Journal of Clinical Endocrinology & Metabolism 86, 930–934 (2001).

27. Hansen, P. S. et al. The relative importance of genetic and environmental factors in the aetiology of thyroid nodularity: A study of healthy Danish twins. Clinical Endocrinology 62, 380–386 (2006).

28. Martin, T. C., Yet, I., Tsai, P.-C. & Bell, J. T. coMET: visualisation of regional epigenome-wide association scan results and DNA co-methylation patterns. BMC Bioinformatics 16, 131 (2015).

29. Moremen, K. W., Tiemeyer, M. & Nairn, A. V. Vertebrate protein glycosylation: diversity, synthesis and function. Nat Rev Mol Cell Biol 13, 448–462 (2012).

30. Li, J. & Ji, L. Adjusting multiple testing in multilocus analyses using the eigenvalues of a correlation matrix. Heredity 95, 221–227 (2005).

31. Huffman, J. E. et al. Polymorphisms in B3GAT1, SLC9A9 and MGAT5 are associated with variation within the human plasma N-glycome of 3533 European adults. Human Molecular Genetics 20, 5000–5011 (2011).

32. Lauc, G. et al. Genomics meets glycomics-the first gwas study of human N-glycome identifies HNF1A as a master regulator of plasma protein fucosylation. PLoS Genet 6, 1–14 (2010).

33. Rijsdijk, V. & Sham, P. C. Analytic approaches to twin data using structural equation models. Briefings in bioinformatics 3, 119–133 (2002).

34. Visscher, P. M. Power of the Classical Twin Design Revisited. Twin Research 7, 505–512 (2004).

35. Medici, M. et al. Identification of novel genetic Loci associated with thyroid peroxidase antibodies and clinical thyroid disease. PLoS Genet 10, e1004123 (2014).

36. Purcell, S. et al. PLINK: A tool set for whole-genome association and population-based linkage analyses. American Journal of Human Genetics 81, 559–575 (2007).

37. Welter, D. et al. The NHGRI GWAS Catalog, a curated resource of SNP-trait associations. Nucleic Acids Res. 42, 1001–1006 (2014).

38. Westra, H. J. et al. Systematic identification of trans eQTLs as putative drivers of known disease associations. Nat Genet 45, 1238–1243 (2013).

39. GTEx Consortium. Human genomics. The Genotype-Tissue Expression (GTEx) pilot analysis: multitissue gene regulation in humans. Science 348, 648–660 (2015).

40. Zeller, T. et al. Genetics and beyond‐‐the transcriptome of human monocytes and disease susceptibility. PLoS ONE 5, e10693 (2010).

41. Buil, A. et al. Gene-gene and gene-environment interactions detected by transcriptome sequence analysis in twins. Nat Genet 47, 88–91 (2015).

42. Drutel, A., Archambeaud, F. & Caron, P. Selenium and the thyroid gland: More good news for clinicians. 78, 155–164 (2013).

43. Schmidt, M. et al. Long-Term Follow-Up of Antithyroid Peroxidase Antibodies in Patients with Chronic Autoimmune Thyroiditis (Hashimoto’s Thyroiditis) Treated with Levothyroxine. Thyroid 18, 755–760 (2008).

44. Kanda, Y. et al. Comparison of biological activity among nonfucosylated therapeutic IgG1 antibodies with three different N-linked Fc oligosaccharides: The high-mannose, hybrid, and complex types. Glycobiology 17, 104–118 (2007).

45. Shields, R. L. et al. Lack of fucose on human IgG1 N-linked oligosaccharide improves binding to human FCγRIII and antibody-dependent cellular toxicity. Journal of Biological Chemistry 277, 26733–26740 (2002).

46. Niwa, R. et al. Enhancement of the antibody-dependent cellular cytotoxicity of low-fucose IgG1 Is independent of FcgammaRNIa functional polymorphism. Clin. Cancer Res. 10, 6248–6255 (2004).

47. Shinkawa, T. et al. The absence of fucose but not the presence of galactose or bisecting N-acetylglucosamine of human IgG1 complex-type oligosaccharides shows the critical role of enhancing antibody-dependent cellular cytotoxicity. Journal of Biological Chemistry 278, 3466–3473 (2003).

48. Niwa, R. et al. Enhanced natural killer cell binding and activation by low-fucose IgG1 antibody results in potent antibody-dependent cellular cytotoxicity induction at lower antigen density. Clinical Cancer Research 11, 2327–2336 (2005).

49. Yamane-Ohnuki, N. et al. Establishment of FUT8 knockout Chinese hamster ovary cells: An ideal host cell line for producing completely defucosylated antibodies with enhanced antibody-dependent cellular cytotoxicity. Biotechnology and Bioengineering 87, 614–622 (2004).

50. Yamane-Ohnuki, N. & Satoh, M. Production of therapeutic antibodies with controlled fucosylation. MAbs 1, 230–236 (2009).

51. Okeley, N. M. et al. Development of orally active inhibitors of protein and cellular fucosylation. Proc. Natl. Acad. Sci. U.S.A. 110, 5404–5409 (2013).

52. Rodien, P. et al. Antibody-Dependent cell-mediated cytotoxicity in autoimmune thyroid disease: relationship to antithyroperoxidase antibodies. Journal of Clinical Endocrinology & Metabolism 81, 2595–2600 (1996).

53. Metcalfe, R. A., Oh, Y. S., Stroud, C., Arnold, K. & Weetman, A. P. Analysis of antibody-dependent cell-mediated cytotoxicity in autoimmune thyroid disease. Autoimmunity 25, 65–72 (1997).

54. Rebuffat, S. A. et al. Human recombinant anti-thyroperoxidase autoantibodies: in vitro cytotoxic activity on papillary thyroid cancer expressing TPO. British journal of cancer 102, 852–861 (2010).

55. Rebuffat, S. A., Nguyen, B., Robert, B., Castex, F. & Peraldi-Roux, S. Antithyroperoxidase antibody-dependent cytotoxicity in autoimmune thyroid disease. The Journal of Clinical Endocrinology & Metabolism 93, 929–934 (2008).

56. Roederer, M. et al. The genetic architecture of the human immune system: A bioresource for autoimmunity and disease pathogenesis. Cell 161, 387–403 (2015).

57. Morris, D. L. et al. Genome-wide association meta-analysis in Chinese and European individuals identifies ten new loci associated with systemic lupus erythematosus. Nat Genet 48, 940–946 (2016).

58. de Lange, K. M. et al. Genome-wide association study implicates immune activation of multiple integrin genes in inflammatory bowel disease. Nat Genet 49, 256–261 (2017).

59. Treviño, L. R. et al. Germline genomic variants associated with childhood acute lymphoblastic leukemia. Nat Genet 41, 1001–1005 (2009).

60. Nutt, S. L., Hodgkin, P. D., Tarlinton, D. M. & Corcoran, L. M. The generation of antibody-secreting plasma cells. Nature Rev Immunology 15, 160–171 (2015).

61. Aust, G. et al. Graves’ disease and Hashimoto’s thyroiditis in monozygotic twins: Case study as well as transcriptomic and immunohistological analysis of thyroid tissues. European Journal of Endocrinology 154, 13–20 (2006).

62. Caturegli, P. et al. Hashimoto’s Thyroiditis: Celebrating the Centennial Through the Lens of the Johns Hopkins Hospital Surgical Pathology Records. Thyroid 23, 142–150 (2013).

63. Armengol, M. P. et al. Thyroid autoimmune disease: demonstration of thyroid antigen-specific B cells and recombination-activating gene expression in chemokine-containing active intrathyroidal germinal centers. The American journal of pathology 159, 861–873 (2001).

64. Weetman, A. P., McGregor, A. M., Lazarus, J. H. & Hall, R. Thyroid antibodies are produced by thyroid-derived lymphocytes. Clinical and experimental immunology 48, 196–200 (1982).

65. Rapoport, B. & McLachlan, S. M. Thyroid autoimmunity. The Journal of clinical investigation 108, 1253–1259 (2001).

66. Beever, K. et al. Highly sensitive assays of autoantibodies to thyroglobulin and to thyroid peroxidase. Clinical Chemistry 35, 1949–1954 (1989).

67. Nakatake, N. et al. Estimation of serum TSH receptor autoantibody concentration and affinity. Thyroid 16, 1077–1084 (2006).

68. Yamauchi, K., Yamada, T., Sato, A., Inazawa, K. & Aizawa, T. Elevation of serum immunoglobulin G in Hashimoto’s thyroiditis and decrease after treatment with L-thyroxine in hypothyroid patients. Internal medicine (Tokyo, Japan) 49, 267–271 (2010).

69. Kawashima, S. T. et al. Serum levels of IgG and IgG4 in Hashimoto thyroiditis. Endocrine 45, 236–243 (2014).

70. Swerdlow, D. I. et al. Selecting instruments for Mendelian randomization in the wake of genome-wide association studies. International journal of epidemiology dyw088 (2016). doi:10.1093/ije/dyw088

71. Haycock, P. C. et al. Best (but oft-forgotten) practices: The design, analysis, and interpretation of Mendelian randomization studies. American Journal of Clinical Nutrition 103, 965–978 (2016).

